# Fentanyl reinforcement history has sex-specific effects on multi-step decision-making

**DOI:** 10.1101/2024.10.10.617707

**Authors:** Eric Garr, Yifeng Cheng, Andy Dong, Patricia H. Janak

## Abstract

It is commonly thought that drug addiction involves a transition to habitual control of action, where the choice to consume drugs becomes automatized and reflects a failure to deliberate over possible negative outcomes. Determining whether the pursuit of addictive drugs is habitual is hampered by a lack of behavior assessments suitable for use during a bout of actual drug seeking. Therefore, to understand how variable histories of drug reinforcement might affect goal-directed and habitual pursuit of drug, we trained rats to perform a multi-step decision-making task to earn oral fentanyl and sucrose rewards following extensive pretraining with either fentanyl or sucrose. Importantly, this task allowed for independent measurements of goal-directed and habitual choice characteristics during online pursuit of rewards, and habitual choice could be further categorized into perseverative and reward-guided components. Chronic fentanyl led to a bias for reward-guided habitual choice specifically in females, and a high degree of perseveration in both sexes. These behavioral changes after chronic fentanyl pretraining generalized across fentanyl and sucrose seeking. In contrast, acute fentanyl selectively increased perseveration in females, and blunted the gradual within-session improvement in goal-directed choice in both sexes. These results show that chronic fentanyl reinforcement promotes habits that generalize across drug and non-drug reward seeking, and that female rats are especially susceptible to habitual control induced by both chronic and acute fentanyl reinforcement.

## Introduction

The drug-seeking behaviors of individuals with substance use disorders are colloquially referred to as “habits”. The term “habit” has acquired the specific definition of an action that is insensitive to its consequences (1). This definition has been applied in drug addiction research, where exposure to addictive drugs has been found to increase habitual reward seeking (2–7). However, this area of research suffers from several limitations. One is that the effect of drugs on habit formation is often measured in the context of seeking nondrug rewards (3–5,8,9). This precludes the study of a drug habit per se. There have been attempts to overcome this limitation by delivering the drug as the instrumental reward on second-order schedules and then attempting to elicit a habit through overtraining (10–12), but this procedure makes it difficult to infer whether responding for drug is outcome-insensitive. Another approach has been to deliver the drug as the instrumental reward and then assess habitual responding by subsequently devaluing the drug reward (2,13) or devaluing the instrumental contingency (14,15), and then testing instrumental performance in extinction. However, an action that is insensitive to devaluation can create ambiguous interpretations. Such an action could be a manifestation of either a “model-free” (MF) mechanism, where actions are driven solely by an expectation of past rewarding outcomes, or perseveration, where action repetition is not based on outcome expectations of any kind (16). These two types of inflexible behavior, possibly orthogonal, cannot be distinguished by devaluation methods. Additionally, the use of reward devaluation makes it difficult to know whether drugs promote habit or impair goal-directed control (17), and this limitation hinders the translational validity of the habit model of drug seeking because it calls into question whether drug-seeking behaviors are habitual (18). Therefore, a different experimental approach is needed.

The two-step task is an alternative method for evaluating goal-directed and habitual components of action control and lends itself to analyses that independently measure these aspects of performance during online reward seeking (19). One strategy that is sometimes used to solve the two-step task, referred to as model-based (MB) learning, requires an agent to represent the probabilities between actions and outcomes such that, when faced with a choice between actions, the subject is able to plan and find the optimal route connecting action with future reward. In contrast, an MF strategy requires an agent only to represent the long-run probabilities of reward associated with each action without accounting for the route connecting action to reward. MB and MF decision-making are often identified with goal-directed and habitual control of action, respectively (20). However, MF choice can vary independently of perseveration in the two-step task, which makes for an additional source of habitual behavior. While MF control of choice is sensitive to prior rewards but insensitive to future action transitions, perseveration is insensitive to both. These features make the two-step task well-suited for addressing how online drug reinforcement alters goal-directed and habitual components of decision-making.

Here, we asked how chronic fentanyl reinforcement affects MB, MF, and perseverative aspects of choice behavior in a two-step task in rats. Previous research has shown that chronic exposure to opioids, including fentanyl, disrupts cognitive flexibility (21–26), but it is less clear whether acute fentanyl exposure changes cognitive flexibility during ongoing decision-making, and what decision processes underlie fentanyl-seeking and how they may be altered by chronic fentanyl. Although there is evidence that opioids bias behavior toward perseveration (27–33), it is unknown whether actions reinforced with opioid reward also show MF features, and whether opioid-induced perseveration is dissociable from these possible changes in MF choice.

We therefore trained rats over weeks to perform a two-step task to earn oral fentanyl or sucrose rewards, followed by a brief period of fentanyl and sucrose sessions in alternation. This experimental design allowed us to dissociate the effects of chronic and acute fentanyl reinforcement on different components of learning and decision-making. We found that chronic fentanyl induced a high degree of perseveration, one measure of habit, in males and females that generalized across fentanyl and sucrose seeking, and also created an MF bias, a second measure of habit, specifically in female rats that also generalized across fentanyl and sucrose seeking. On the other hand, acute fentanyl induced within-session changes in decision-making; acute fentanyl interfered with a progressive within-session improvement in MB choice and also increased perseveration in females given brief, but not extensive, fentanyl training.

## Materials and Methods

All procedures were approved by the Johns Hopkins University Animal Care and Use Committee. To examine the reinforcing properties of oral fentanyl, Long-Evans rats (4 males, 4 females) were water-restricted and trained to press a left and right lever on a fixed-ratio 1 schedule for oral fentanyl (fentanyl hydrochloride dissolved in tap water, Cayman Chemical Company; 50 µg/ml, 0.05 ml per delivery) over four daily 1-hour sessions (two per lever). Each rat was then given the opportunity to press one lever for fentanyl and the other lever for tap water while thirsty or not thirsty. Lever outcome, testing order, and sex were counterbalanced. Rats were then placed back on water restriction and trained to press one lever on a fixed-ratio 5 schedule for oral fentanyl (50 µg/ml, 0.05 ml per delivery) and another lever for sucrose (10% solution in tap water, 0.05 ml per delivery) over two daily 1-hour sessions (1 per lever). Each rat was then given the opportunity to press each lever for their respective outcomes 30 minutes after IP injections of naltrexone (0.1 mg/kg) or saline in separate 45-minute sessions. Lever outcome, testing order, and sex were counterbalanced. Rats were given 1 hour of supplemental water after each session. During all behavioral sessions, only one lever was available to press at a time.

To assess behavioral strategy during pursuit of fentanyl, a separate cohort of thirsty Long-Evans rats was trained to perform a two-step task to earn oral fentanyl (25 µg/ml; 9 males and 8 females) or 10% sucrose (10 males and 8 females). The fentanyl concentration was set to 25 µg/ml to increase the number of trials the rats were willing to perform. Following shaping (see *Supplemental Materials*), training on the full task commenced and continued for 22-30 daily sessions. Each trial began with illumination of the center magazine, prompting the rat to initiate the trial with a center nose-poke, followed by a choice between the left and right nose-poke ports. This choice triggered the insertion of one of the two levers on the opposite wall. Each nose port was predominantly associated with different levers, with transition probabilities fixed at 0.8 and 0.2 (common and rare transitions, respectively). Upon lever-pressing, reward was delivered with a probability of 0.8 if the rat pressed the high-reward lever, and 0.2 if the rat pressed the low-reward lever. A clicker sound accompanied reward delivery, while 1-s white noise accompanied reward omission. The reward probability associated with each lever was randomly alternated after a minimum of 20 trials, with a 10% chance of switching on each subsequent trial, and a maximum of 35 trials before a switch occurred. Reward size was fixed at 0.05 ml. During each session, the fentanyl group was free to earn up to 150 rewards, while the sucrose group was limited to the average number earned by the fentanyl group during the previous session. This was done to equate the average number of rewards between groups.

Following training on the two-step task, all rats received alternating sessions with fentanyl and sucrose rewards (6 sessions per reward, 12 total). Sessions with the unfamiliar reward occurred in an altered context with lemon scent and honeycomb textured floors. The maximum number of sucrose rewards per session per rat was set to the number of fentanyl rewards earned during the previous session.

## Results

### Pharmacological properties of oral fentanyl drive intake

We first confirmed that oral fentanyl is reinforcing beyond its thirst-quenching properties by training rats to press a lever for either oral fentanyl or water during conditions of thirst and non-thirst (**Fig 1A**). Rats escalated within-session intake of fentanyl more quickly than water regardless of deprivation state (main effect of reward: *F*(1,7) = 9.80, *p* = .017; no reward x deprivation interaction: *F*(1,7) = .031, *p* = .866). We additionally confirmed that instrumental responding for oral fentanyl depends on activation of opioid receptors (**Fig 1B**). The opioid antagonist, naltrexone, significantly affected pressing for fentanyl, but not sucrose (reward x injection interaction: *F*(1,7) = 7.05, *p* = .033). Rats took longer to terminate pressing for fentanyl after naltrexone relative to saline, most likely because the drug intake threshold was increased following opioid receptor blockade (**Fig 1B**). These results show that oral fentanyl motivates instrumental responding in a way that depends on opioid receptor activation.

**Figure 1.**
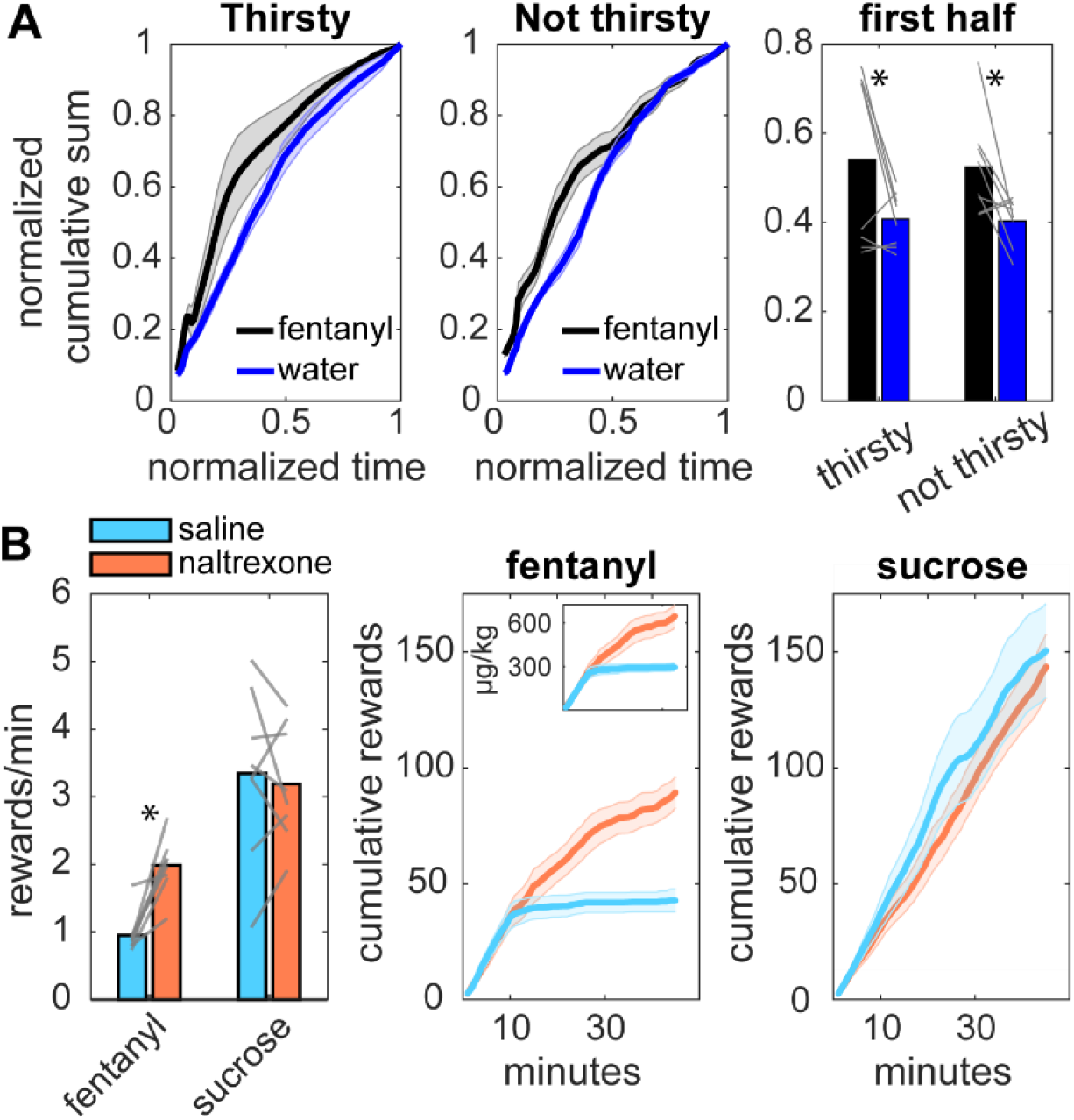
**(A)** Cumulative sum of fentanyl and water rewards as a function of session time as rats pressed a lever for liquid rewards under thirsty and non-thirsty conditions. Cumulative sums were normalized in the range of 0-1 and then interpolated. Right-most bar plot shows means over the first half of the session. **(B)** Left: Mean reward rates while rats pressed a lever on an FR5 schedule for either liquid fentanyl or sucrose after receiving IP injections of saline or naltrexone. Grey lines are individual rats. Middle: Cumulative fentanyl rewards earned across sessions. Inset shows the cumulative intake. Right: Cumulative sucrose rewards earned across sessions.

### In females, extensive training with fentanyl biases behavior towards model-free choice

To understand how fentanyl reinforcement history affects decision-making, we trained two new groups of rats to earn either oral fentanyl or sucrose rewards on a two-step task (**Fig 2A**; 34–39). The training period lasted 22-30 sessions, ensuring rats fully acquired knowledge of the task structure. Training was followed by the opportunity for all rats to earn either fentanyl or sucrose in alternating sessions (6 test sessions per reward type; **Fig 2B**). This experimental design allowed us to investigate how short-term vs long-term drug reinforcement histories impact decision-making for drug and natural rewards. Analyses were conducted on the final six sessions of alternating rewards (i.e. 3 sessions per reward type), unless otherwise noted.

**Figure 2.**
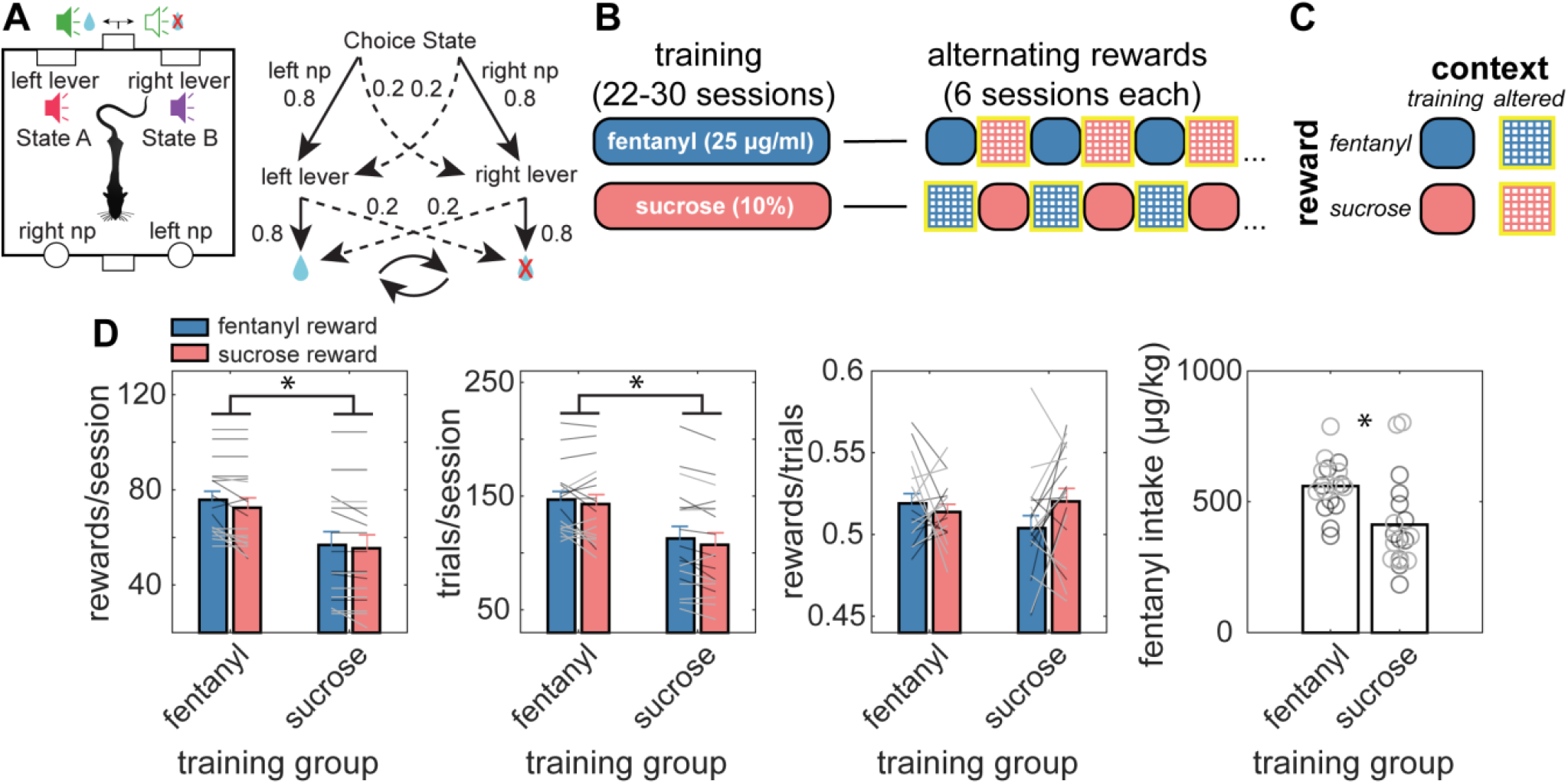
Two-step task structure, experimental design, and performance statistics. **(A)** Left: Schematic of behavioral apparatus, consisting of one magazine for center fixation and two flanking nose-ports, and one magazine for reward and two retractable levers on the opposite side. Right: task structure consisted of an initial choice state where either the left or right nose-poke could be chosen, and each choice led to a common (0.8) or rare (0.2) lever transition probability. Each lever was associated with a high (0.8) or low (0.2) reward probability, which switched in blocks. **(B)** Illustration of experimental design. Two groups of rats were trained to earn either oral fentanyl or sucrose on the two-step task for many sessions, and then experienced alternating sessions of fentanyl and sucrose. Alternating sessions with the reward type not experienced during training took place in contexts with a different scent and floor texture. **(C)** Definition of symbols in panel A. During the alternating rewards phase, two variables changed between sessions: reward type and context. The training context refers to the context the rats were trained in, while the alternating context refers to an altered floor texture (indicated by grid pattern) and scent (indicated by yellow outline). **(D)** Mean rewards, trials, reward rate, and fentanyl intake per session. Data are taken from the final 6 sessions of the alternating rewards phase. Black data points are males and grey data points are females.

We first analyzed basic performance statistics from these fentanyl and sucrose sessions (**Fig 2D**). The number of rewards earned and trials performed were not affected by the instrumental reward (no main effects or interactions: *F*’s(1,31) < 1.52, *p*’s > .23), but were significantly affected by group (main effect: *F*’s(1,31) > 8.94, *p*’s < .005) and sex (main effect: *F*’s(1,31) > 10.65, *p*’s < .003). The fentanyl training group completed more trials and earned more rewards than the sucrose training group, and males outperformed females. However, when examining fentanyl intake (**Fig 2D**), the effect of sex went away (*F*(1,31) = 2.50, *p* = .12) but the effect of group remained (*F*(1,31) = 9.31, *p* = .005). These results show that rats with an extensive history of fentanyl performed more trials and earned more rewards, but this was likely a side effect of how total rewards were controlled during test sessions (i.e. fentanyl rewards are practically unlimited and determine the limit for sucrose rewards, and the sucrose training group was less familiar with fentanyl).

To examine how fentanyl reinforcement history affects the balance between MF and MB choice, we next analyzed session-averaged stay probabilities accounting for previous trial outcome and transition, as well as instrumental reward, training group, and sex (**Fig 3A**). The stay probability is the probability of repeating the choice from the previous trial. A purely MF agent will show high stay probabilities following rewarded trials compared to omission trials regardless of whether the choice-lever transition is common or rare, while a purely MB agent will consider how likely each choice will transition to each lever. An MF index was calculated from the stay probabilities as (reward-common + reward-rare) – (omission-common + omission-rare), and an MB index was calculated as (reward-common + omission-rare) – (reward-rare +omission-common).

**Figure 3.**
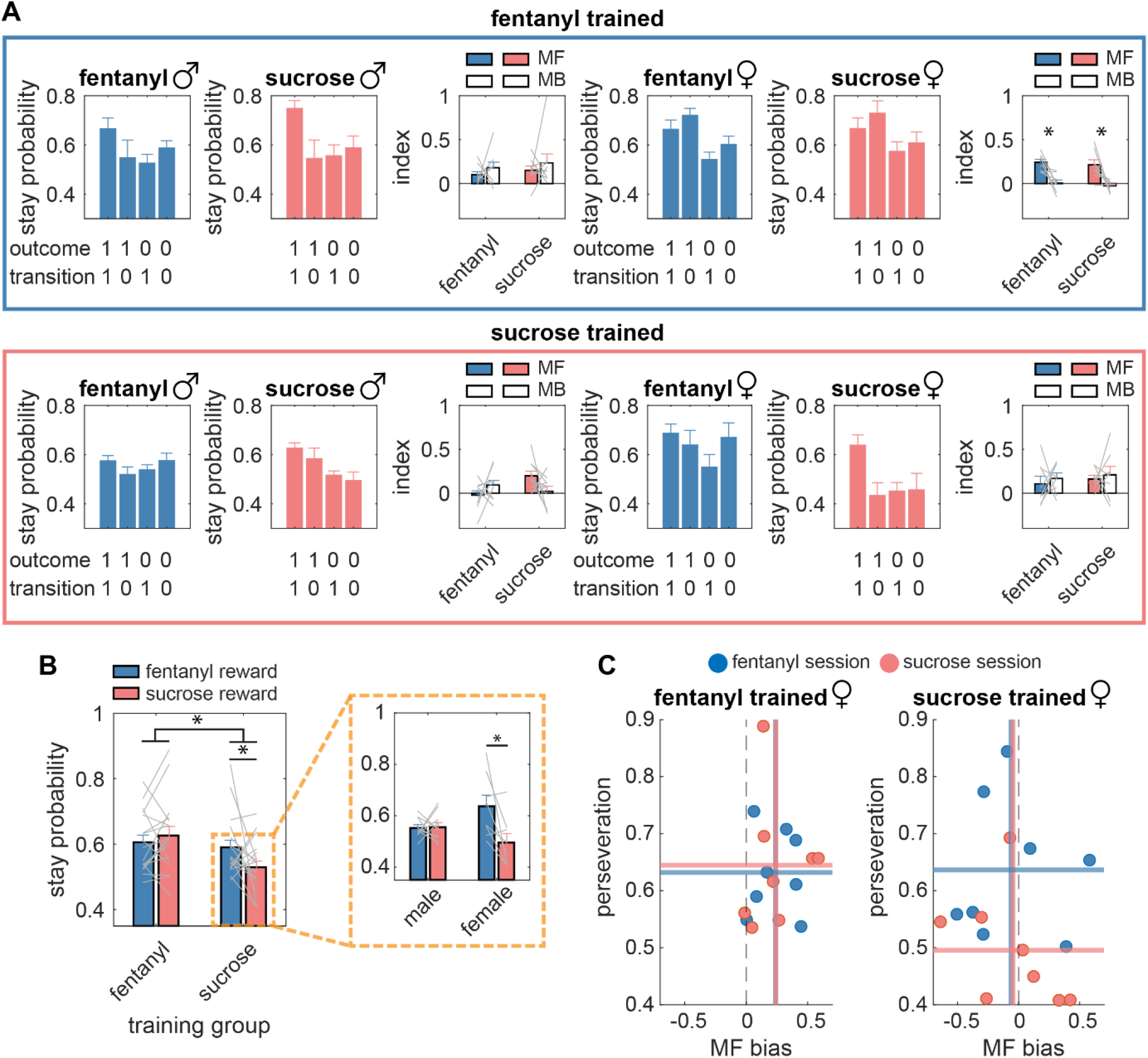
Fentanyl enhances habit expression in female rats. **(A)** Stay probabilities plotted as a function of the previous trial outcome (1 = reward, 0 = omission), transition (1 = common, 0 = rare), training group (fentanyl vs. sucrose), instrumental reward (fentanyl vs. sucrose), and sex (male vs. female). MF and MB indices derived from the stay probabilities are also shown. Bars show means across rats, error bars show SEMs, and grey lines are individual rats. **(B)** Stay probabilities averaged over trial-level variables (i.e. outcome and transition) and plotted as a function of training group and instrumental reward (left). Data are broken down further by dividing the sucrose training group into males and females (right). Bars show means across rats, error bars show SEMs, and grey lines are individual rats. **(C)** Summary of main findings. Perseveration, defined as the mean stay probability, is plotted against MF bias, defined as MF – MB index. Fentanyl trained females (left) showed a high level of perseveration and a high MF bias during fentanyl and sucrose sessions. Sucrose trained females (right) showed a high level of perseveration specifically during fentanyl sessions, but not MF bias. Each data point is an individual rat.

Rats showed greater stay probabilities following reward vs. omission trials (main effect of outcome: *F*(1,31) = 55.28, *p* < .001), which is consistent with MF choice. In addition, stay probabilities were affected by transition type (outcome x transition interaction: (*F*(1,31) = 14.70, *p* < .001), resulting in an MB pattern of stay probabilities: a greater stay probability after reward-common vs. reward-rare trials, and a greater stay probability after omission-rare vs. omission-common trials (simple main effects, *p*’s < .011). These results show that rats’ choices were a mixture of MF and MB strategies. Critically, we found that the extent to which rats relied on MF and MB strategies depended on fentanyl history and sex. Females given extensive prior training with fentanyl showed a pattern of stay probabilities consistent with a low degree of MB choice (outcome x transition x group x sex: *F*(1,31) = 9.21, *p* = .005). An analysis of MF and MB indices showed a significant index x group x sex interaction (*F*(1,31) = 6.13, *p* = .019), with a significantly higher MF bias (MF index – MB index) in females given extensive training with fentanyl (*p* = .009; **Fig 3A**). Notably, MF bias could not be predicted from the number of trials performed (*β*’s < -.002, *p*’s > .143). We also applied the same analyses to multi-trial regression coefficients, and found similar results (**Fig S1**). These analyses show that, overall, there was a mixture of MF and MB choice across rats, but in females given extensive training with fentanyl, MF indices were high while MB indices were greatly decreased.

### Female rats are more susceptible to the acute effects of fentanyl on perseveration

When stay probabilities are averaged over trial-level variables, they measure perseveration—the overall tendency to repeat actions. Our analysis uncovered significant effects of fentanyl on perseveration. Perseveration was greater for rats trained with fentanyl than sucrose (main effect of group: *F*(1,31) = 4.75, *p* = .037; **Fig 3B**), and acute fentanyl reinforcement increased perseveration even for rats trained with sucrose (reward x group interaction: *F*(1,31) = 5.98, *p* = .02; **Fig 3B**). To investigate whether the acute effect of fentanyl on perseveration was sex-dependent, stay probabilities from the sucrose training group were divided by sex (**Fig 3B**). Females, but not males, showed a greater tendency to perseverate during fentanyl versus sucrose sessions (sucrose training group, sex x reward interaction: *F*(1,16) = 6.80, *p* = .019); **Fig S2**).

These results show that extensive training with fentanyl increased perseveration in a way that generalized across reward identity and sex, while the perseveration observed within fentanyl-reinforced sessions after extensive sucrose training was highly specific to reward identity (fentanyl) and sex (females). Perseveration could not be predicted by the number of trials performed (*β*’s < -.001, *p*’s > .051). Interestingly, there were no significant correlations between perseveration and MF bias in females (**Fig 3C**; -.46 < *r*’s < .12, .25 < *p*’s < .99), supporting the idea that extensive and brief fentanyl reinforcement histories have distinct behavioral effects in female rats. We also found no correlations between perseveration and MF or MB indices across sexes (**Fig S3)**, implying that perseveration is a separate dimension of choice.

It could be argued that these analyses do not precisely assess acute effects of fentanyl in isolation from chronic effects because we ignored data from the first few sessions when rats were initially exposed to fentanyl. We chose to ignore these sessions because rats were becoming familiar with the new rewards and altered contexts, which could interfere with performance. Nevertheless, we examined the first fentanyl and sucrose sessions for rats given prior training with sucrose to uncover whether acute fentanyl altered perseveration during the first exposure to the drug. Indeed, perseveration was greater during the first fentanyl session compared to the subsequent sucrose session, although there was no interaction with sex (**Fig S4**). On average, perseveration increased over the course of the fentanyl session and decreased during the sucrose session (**Fig S4**), although this difference was not evident in a reward x session half interaction (*F*(1,16) = 2.28, *p* = .150). However, the degree of perseveration during fentanyl sessions was positively correlated with the number of fentanyl rewards earned (*r* = .47, *p* = .048). This was not true of sucrose (*r* = .13, *p* = .62). This analysis shows that the effect of fentanyl on perseveration was evident from the first encounter with fentanyl, and that this is likely an acute pharmacological effect.

### Acute fentanyl reinforcement stifles within-session improvement in model-based choice

When rats earned fentanyl rewards on the two-step task, they accumulated fentanyl over the course of a session. Therefore, session-averaged measures may not provide a full description of how fentanyl affects choice. We analyzed within-session changes in stay probabilities by splitting the sessions into halves (**Fig 4A**). MB choice grew in strength during sucrose sessions, but showed a decline during fentanyl sessions. A mixed model ANOVA revealed an outcome x transition x reward x half interaction (*F*(3,93) = 4.27, *p* = .047), and simple main effects tests showed that, during the second half of the session, stay probabilities following reward-common trials were smaller during fentanyl versus sucrose sessions (*p* = .006), while stay probabilities following reward-rare trials were greater during fentanyl versus sucrose sessions (*p* = .031).

**Figure 4.**
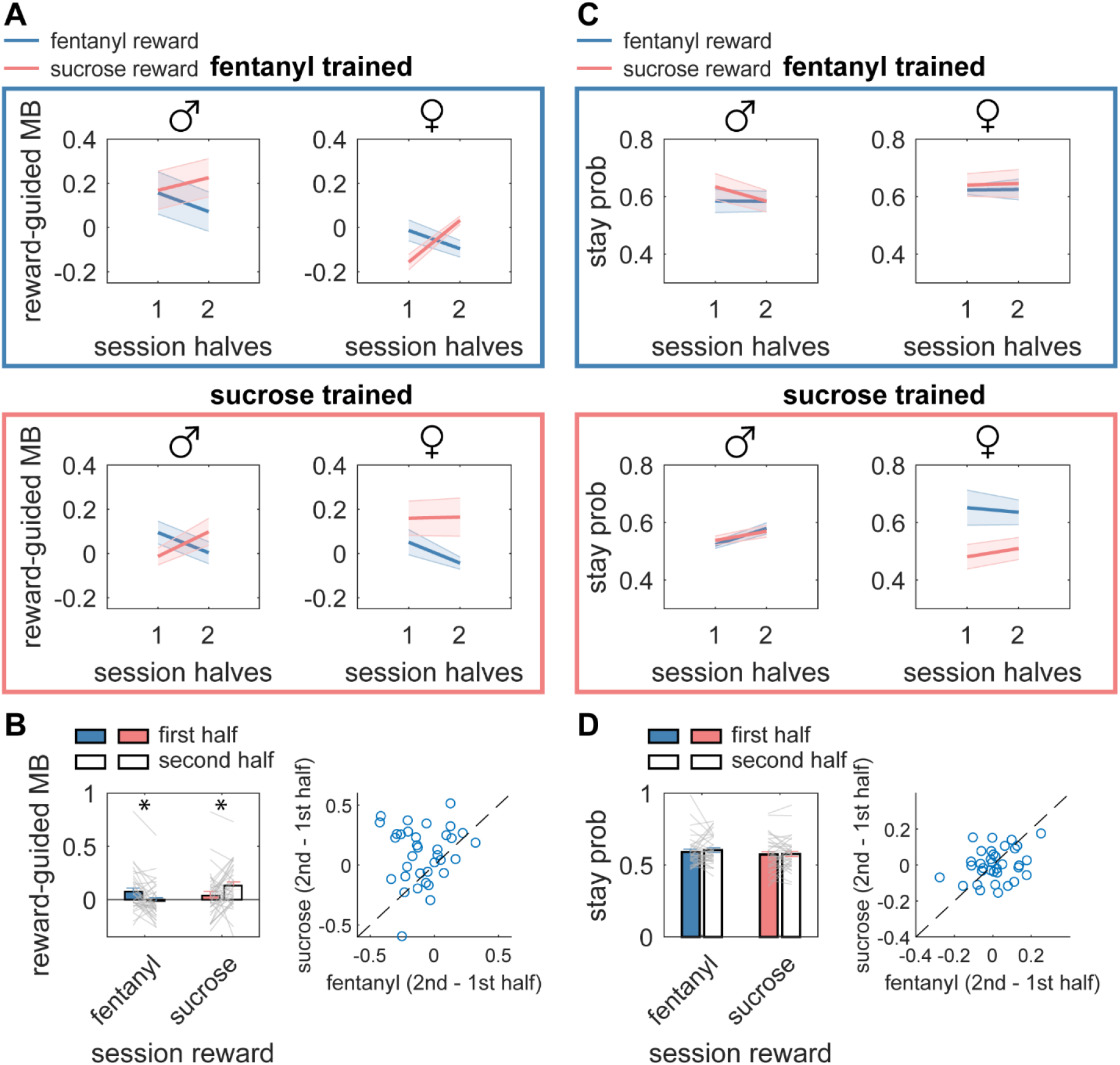
Within-session decrements in MB choice during fentanyl reinforcement. **(A)** Reward-guided MB indices, defined as the difference in the stay probabilities between reward-common and reward-rare trials, are shown as a function of session half separately for males and females trained with fentanyl (top) or sucrose (bottom). Lines are means across rats, and shaded bounds are SEMs. **(B)** Summary of main finding. Reward-guided MB indices are plotted as a function of instrumental reward and session half for all rats (left). The difference between reward-guided MB choice during the first and second halves of the sessions is plotted for individual rats during sucrose and fentanyl sessions. **(C)** Stay probabilities averaged across trial-level variables (i.e. outcome and transition) are plotted over sessions halves separately for instrumental reward, training group, and sex. **(D)** Summary of perseveration scores over time. Stay probabilities are plotted as a function of instrumental reward and session half for all rats (left). The difference between perseveration scores during the first and second halves of the sessions is plotted for individual rats during sucrose and fentanyl sessions.

To visualize this effect, we computed reward-guided MB indices by subtracting the stay probabilities between reward-common trials and reward-rare trials. Across training groups and sexes, there was a clear increase in reward-guided MB choice over the course of sucrose sessions, while this behavior decreased during fentanyl sessions (**Fig 4B**). Notably, session halves were divided by trials, not time. This means that each session half should contain roughly equal numbers of rewards, indicating that time-dependent differences in MB choice for fentanyl versus sucrose are not due to differences in reward rate or motivation. Indeed, there was no difference in rewards earned between session halves (*F*(1,34) = 2.49, *p* = .12). Perseveration did not show changes across these sessions (**Fig 4C and D**; no main effect or interactions with session half, *p*’s > .143). Thus, a component of model-based choice for sucrose reward showed a progressive improvement within sessions—an effect not observed for acute fentanyl. This trend generalized across training groups and sexes, suggesting the decline in MB choice under fentanyl reflects an acute effect that is not dependent on prior chronic fentanyl.

## Discussion

We find that fentanyl-seeking actions are driven by different combinations of MB, MF, and perseverative features depending on the history of fentanyl reinforcement and biological sex. Choices that led to fentanyl were largely MF in female rats only after extensive prior training with fentanyl. However, this type of habitual behavior also extended to sucrose seeking, which suggests that this is not a “drug habit” but rather a general change in decision-making. The finding that most resembled a drug-selective habit was the within-session decrement in MB choice induced by acute cumulative fentanyl intake across sexes and training groups. Yet given the alignment between MB choice and goal-directed action selection, it would be more accurate to state that acute fentanyl impaired goal-directed control rather than inducing a change in habits. We also found that MF choice was largely independent of perseveration, both in terms of how the variables map onto orthogonal dimensions and the drug reinforcement conditions that give rise to their changes. Female rats given brief training with fentanyl showed an acute increase in perseveration during fentanyl relative to sucrose sessions, and this change in perseveration was not accompanied by changes in MF choice. This work demonstrates that drug-induced habits can arise from either an MF or perseverative mechanism, depending on drug reinforcement history and sex. Together, these findings provide new insight into the impacts of acute and chronic fentanyl on decision making processes in males and females.

To understand how chronic and acute fentanyl reinforcement affect the balance between MF and MB choice, we trained rats to earn oral fentanyl and sucrose rewards in separate sessions after extensive training on either fentanyl or sucrose. We chose to deliver fentanyl as an oral reward to compare with a control oral sucrose condition. While the most common preclinical model of fentanyl seeking is intravenous self-administration, oral delivery of fentanyl reinforces an instrumental action in a way that mirrors behavioral effects of intravenous fentanyl: inelastic demand curves, escalation of intake, and cued reinstatement (41), and is a common route of administration for humans. We show the reinforcing properties of oral fentanyl in thirsty rats at least partially depend on activation of opioid receptors, suggesting that reward-seeking is driven by pharmacological effects of fentanyl itself.

Using a two-step task, we showed that decision-making processes in male and female rats are affected differently by chronic and acute fentanyl reinforcement. While chronic fentanyl reinforcement led to a high level of perseveration that generalized across sexes, the same drug history led to a bias toward MF choice for both fentanyl and sucrose only in female rats. There is some evidence in the literature that females respond differently to chronic fentanyl compared to males. After many repeated fentanyl exposures, female rodents self-administer fentanyl at higher rates, are more prone to fentanyl relapse, and show greater resistance to extinction of fentanyl seeking (41–47). The basis of these sex differences may lie in neuroendocrinology (48). However, the evidence in favor of higher susceptibility to opioid craving and relapse in females is mixed (49), and in humans, it has been found that men overdose at higher rates than women (50). The present study extends this work to show sex-specific effects of chronic fentanyl-induced changes in action control, which, notably, cannot be attributed to differences in drug intake.

The behavioral changes after chronic fentanyl contrast with those observed in rats given a relatively brief number of sessions with fentanyl following extensive sucrose training. Females showed an acute increase in perseveration during fentanyl-seeking relative to sucrose-seeking, despite displaying balanced MF and MB choice. This acute fentanyl-induced perseveration is probably a result of bioactive fentanyl, as perseveration was also high during the very first session with fentanyl. Another manifestation of acute fentanyl was that, across sexes and in both chronic fentanyl and sucrose training groups, the accumulation of fentanyl within a session led to a decrease in MB control. In contrast, MB control gradually improved during sucrose seeking. An acute disruption in behavioral flexibility may increase the risk of developing future drug abuse, as compromised cognitive flexibility is associated with greater vulnerability to addiction and predicts future escalation of drug use (22,36,51,52). However, it is uncertain whether compromised cognitive flexibility causes addictive behaviors or whether they are symptoms of the same underlying cause.

In summary, we found that rats can perform a multi-step decision-making task that yields measures of model-based, model-free, and perseverative choice characteristics that are modulated by fentanyl reinforcement history. It would not be possible to unearth these behavioral differences with the standard methodology of measuring responding under free-operant schedules, highlighting the utility of combining the two-step task with drug reward for future directions in drug habit research. The finding that the length of prior fentanyl use and biological sex interact to produce distinct behavioral deficits may help to explain why some individuals transition from recreational drug use to addiction. The current set of results would be strengthened by extending the experimental design to include a non-water-deprived condition, incorporating fentanyl self-administration via the common intravenous route, and studying how distinct features of action control map onto and predict different features of addiction.

## Author contributions

EG and PHJ designed the experiments. EG and AD collected the data. EG visualized and analyzed the data with input from YC and PHJ. EG and PHJ prepared the manuscript with input from YC. Data and code are available upon publication and request to.

