## Supplemental figure for "Fentanyl reinforcement history has sex-specific effects on multi-step decision-making"

### SUPPLEMENTAL MATERIAL

#### Animals

Long-Evans rats were purchased from Envigo. Rats were kept in a temperature- and humidity-controlled environment with a light:dark cycle of 12:12 h (lights on at 7:00 a.m.). Experiments were conducted during the light cycle. All animal care and experimental procedures were approved by the Johns Hopkins University Animal Care and Use Committee.

#### Behavioral Training

##### *Two-step task shaping*

Shaping occurred in four stages.

1. Magazine training: On the first day, rats were allowed to freely explore a standard behavioral chamber (Med Associates) and collect 30 non-contingent rewards (0.1 ml per delivery of solution) delivered randomly at an average interval of 60 seconds to a reward magazine centered on a side wall.
2. Lever press training: Rats were then trained to press the levers flanking the reward magazine under a fixed-ratio 1 (FR1) schedule until they earned a total of 50 rewards (0.05 ml per reward). Reward was always accompanied by a clicker sound. Only one lever was available at a time.
3. Lever trigger training: Lever insertion was then made contingent on nose-poking. Rats were trained to nose-poke into simultaneously illuminated left and right ports on the opposite wall, triggering the insertion of a single lever on the other side of the chamber. This action triggered a distinct tone: a high pitch indicated the insertion of the left lever, while a low pitch indicated the right lever. Each port was predominantly associated with one lever, with transition probabilities fixed at 0.8/0.2. Pressing the available lever resulted in reward with 100% certainty. Rats were free to choose which port to poke in on each trial.
4. Nose port trigger training: Nose port illumination was then made contingent on poking into a center magazine. Each trial began with illumination of the center magazine placed between the left and right nose ports. Upon entry into this magazine, the trial proceeded as described in step 3.

#### Data analysis

##### *Stay probability analysis*

To assess decision-making strategies, we analyzed the probability of rats repeating their previous first-step choice ("stay probability") based on the outcome and transition of the previous trial. Trials were classified into four conditions: (1) common transition with reward, (2) common transition with reward omission, (3) rare transition with reward, and (4) rare transition with reward omission. Stay probability was calculated as the proportion of trials where the rat repeated the same first-step choice as in the previous trial. Stay probability analyses were conducted on all trials.

To calculate the model-based (MB) and model-free (MF) indices, we used the following formulas:

Model-based (MB) index: This index was calculated as the sum of the probability of staying on the same choice following reward-common trials (RC) and omission-rare trials (OR), minus the sum of the probability of staying on the same choice following reward-rare trials (RR) and omission-common trials (OC). Mathematically, this is represented as:

$$MB = [p(stay|RC) + p(stay|OR)] - [p(stay|RR) + p(OC)]$$

Model-free (MF) index**:** This score was calculated as the sum of the probability of staying on the same choice following a reward, regardless of the transition type, minus the sum of the probability of staying on the same choice following no reward, regardless of the transition type. Mathematically, this is represented as:

$$MF = [p(stay|RC) + p(stay|RR)] - [p(stay|OC) + p(stay|OR)]$$

***Multi-trial-back regression analysis***

To assess the influence of past transitions and outcomes on future decisions, we conducted a logistic regression analysis that considers the effects of multiple previous trials (Miller et al., 2016; Miller et al., 2017). A trial was coded as +1 if the left port was chosen, and -1 if the right port was chosen. For each of the four possible trial types (RC, RR, OC, OR), binary vectors were created to indicate whether each trial belonged to a given trial type (+1 for trials of a given type where the rat selected the left port, an -1 for trials of a given type where the rat selected the right port). The following regression model was used:

$$\log\left(\frac{p_{left}(t)}{p_{right}(t)}\right) = \sum_{a=1}^{T} \beta_{RC}(\tau) * RC(t-\tau) + \sum_{a=1}^{T} \beta_{RR}(\tau) * RR(t-\tau) + \sum_{a=1}^{T} \beta_{OC}(\tau) * OC(t-\tau) + \sum_{a=1}^{T} \beta_{OR}(\tau) * OR(t-\tau)$$

In this model, the regression coefficients $\beta_{RC}$, $\beta_{RR}$, $\beta_{OC}$, and $\beta_{OR}$ capture the likelihood of repeating a choice made $\tau$ trials ago, contingent on the specific outcome and transition type, while $T$ represents the number of past trials considered by the model for predicting future choices. To compute MB and MF indices that account for influences from multiple past trials, we summed the relevant regression coefficients:

$$MB = \sum_{a=1}^{T} [\beta_{RC}(a) + \beta_{OR}(a)] - \sum_{a=1}^{T} [\beta_{RR}(a) + \beta_{OC}(a)]$$

$$MF = \sum_{a=1}^{T} [\beta_{RC}(a) + \beta_{RR}(a)] - \sum_{a=1}^{T} [\beta_{OC}(a) + \beta_{OR}(a)]$$

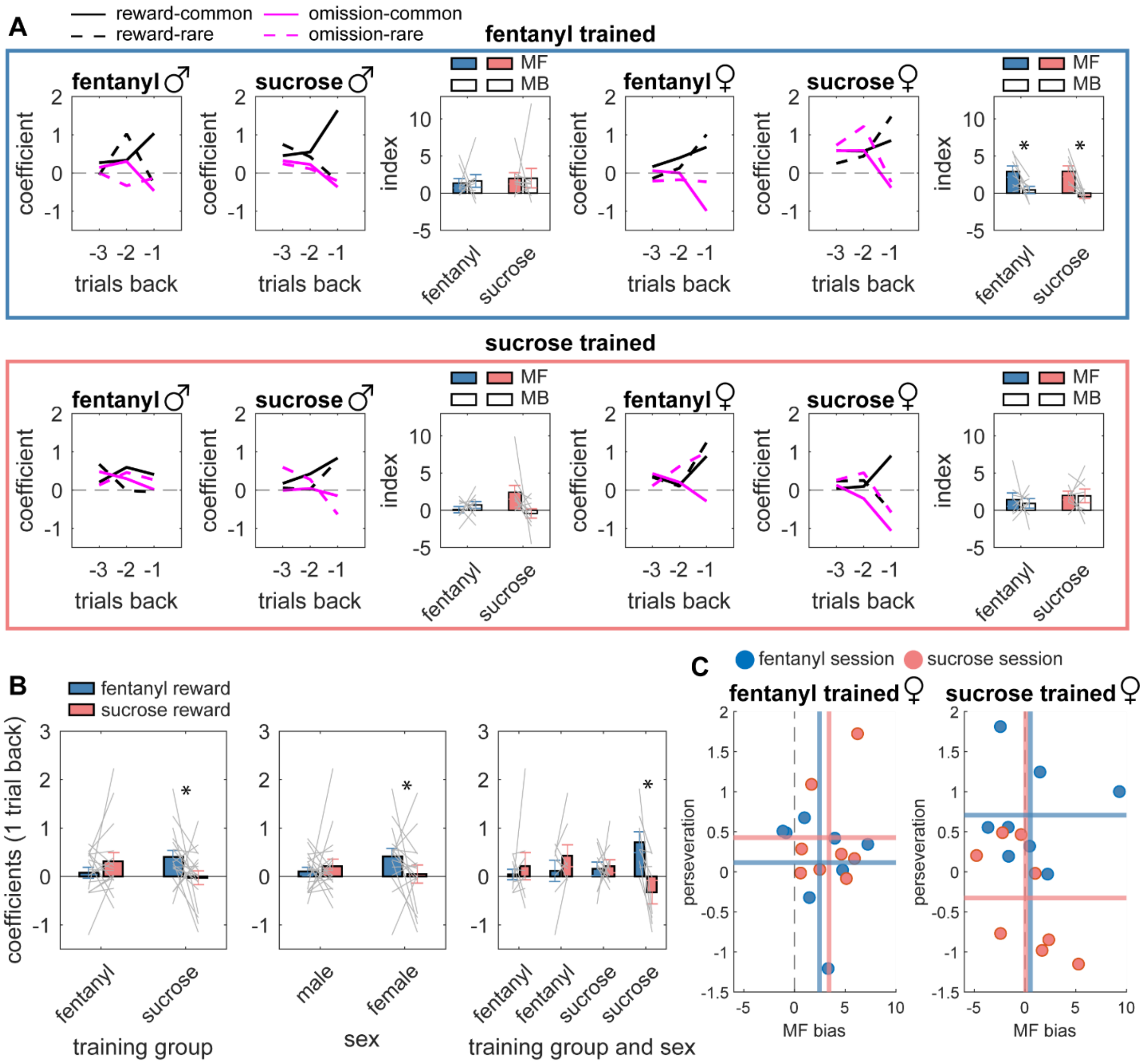


**Figure S1. (A)** Regression coefficients plotted as a function of trial outcome and transition up to three trials in the past, show separately for training group, instrumental reward, and sex. The coefficients capture the likelihood of repeating a choice made up to three trials in the past (i.e. a positive coefficient means high likelihood of repeating choice and a negative coefficient means high likelihood of switching choice). Like the stay probability analysis (**Fig 3A**), females given extensive prior training with fentanyl showed coefficients consistent with a low degree of MB choice, but only for 1 trial back in time (outcome x transition x group x sex x trials, $F(2,62) = 5.36$, $p = .007$). Therefore, MF and MB indices were computed from coefficients only one trial back. Lines and bars show means across rats, error bars show SEMs, and grey lines are individual rats. **(B)** Regression coefficients one trial back averaged over trial-level variables (i.e. outcome and transition) and plotted as a function of training group and instrumental reward (left), sex and instrumental reward (middle), and all three variables combined (right). Like the stay probability analysis (**Fig 3B**), females, but not males, showed a greater tendency to perseverate during fentanyl versus sucrose sessions (reward x group x sex: $F(1,31) = 9.11$, $p =$

.005). Bars show means across rats, error bars show SEMs, and grey lines are individual rats. **(C)** Perseveration, defined as the mean regression coefficient one trial back, is plotted against MF bias, defined as MF – MB index. Each data point is an individual rat.

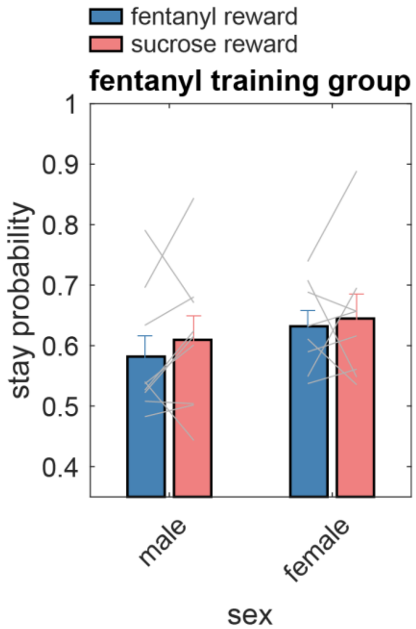

**Figure S2.** Mean stay probabilities for males and females within the fentanyl training group.

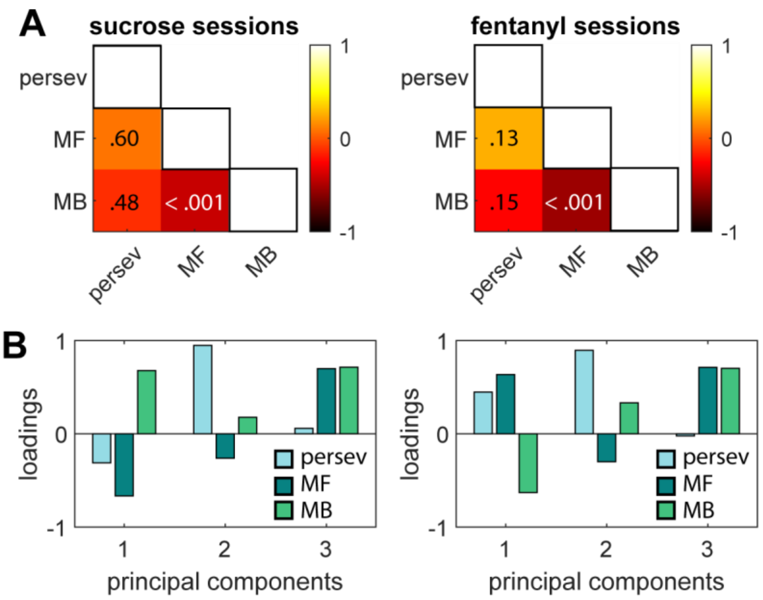


**Figure S3. (A)** Heat map showing correlations between perseveration scores, MF indices, and MB indices separately for sucrose and fentanyl sessions. Colors represent correlation coefficients, and text inside matrix cells are *p* values. **(B)** Principle component analysis on perseveration scores, MF indices, and MB indices separately for sucrose and fentanyl sessions. Perseveration loads overwhelmingly on PC2.

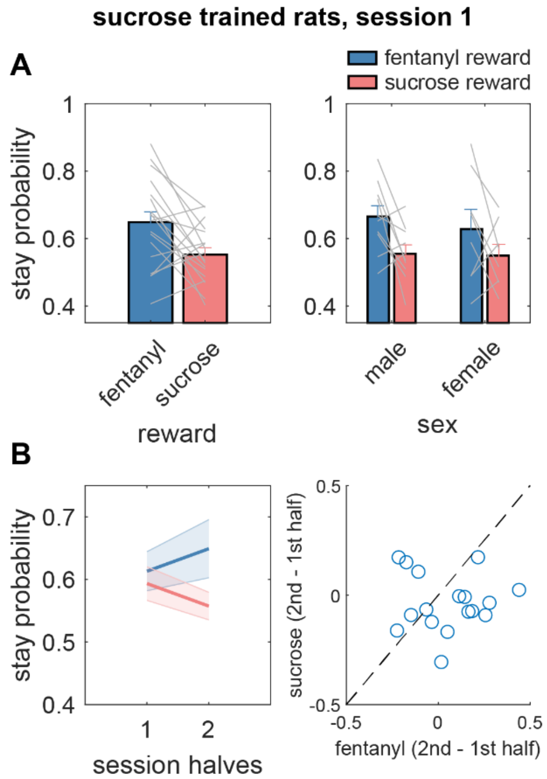


**Figure S4. (A)** Mean stay probabilities from the first fentanyl session, and the subsequent sucrose session, for rats previously given extensive training with sucrose. Data are shown collapsed across (left) and divided by (right) sex. **(B)** Mean stay probabilities divided by session halves and reward type (left). The difference between halves is shown for individual rats for fentanyl and sucrose sessions (right).
